## Supplementary figures and images for "Leveraging a founder population to identify novel rare-population genetic determinants of lipidome"

### Supplementary Figure 1

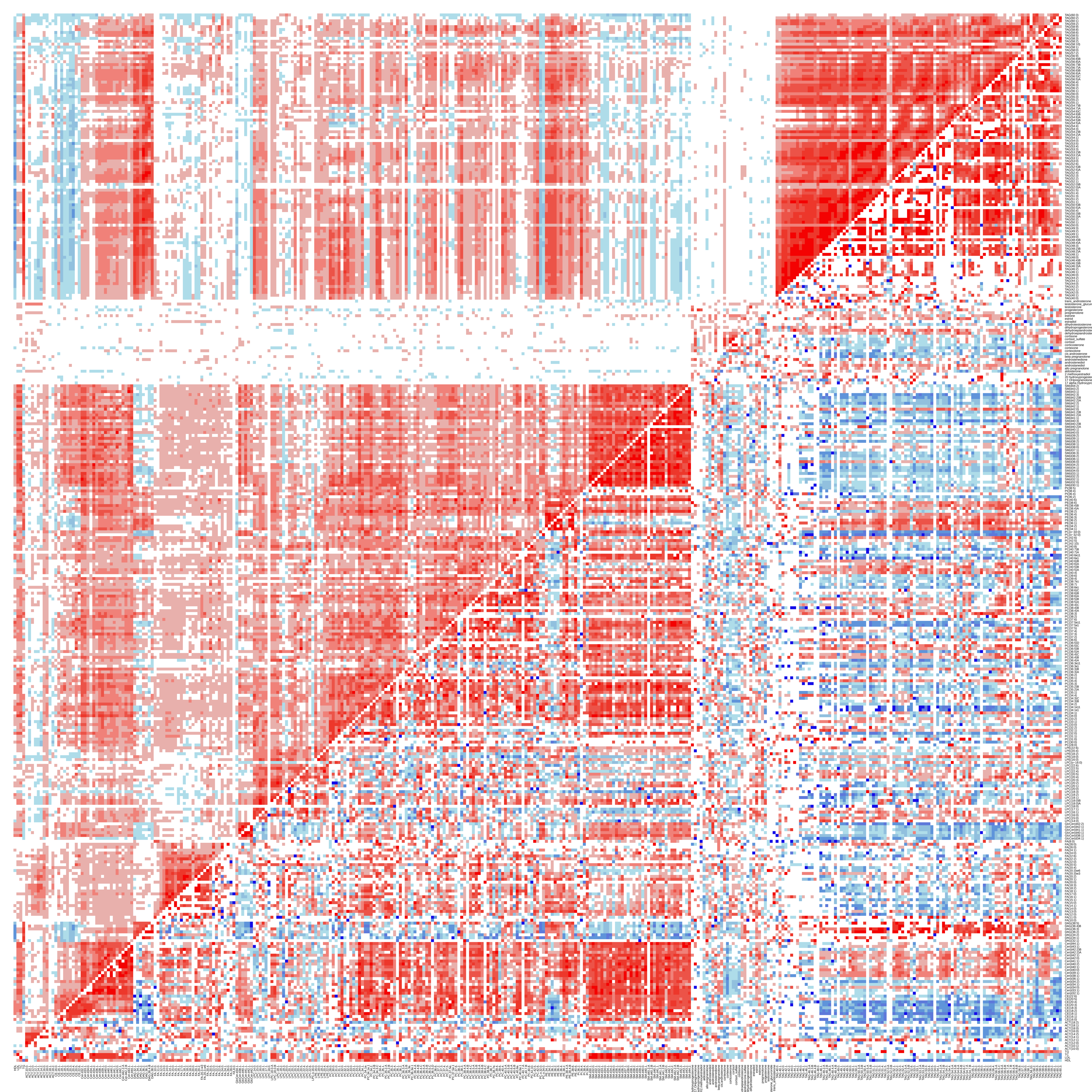
